# A high-sensitivity low-nanoflow LC-MS configuration for high-throughput sample-limited proteomics

**DOI:** 10.1101/2023.04.27.538542

**Authors:** Runsheng Zheng, Manuel Matzinger, Rupert Mayer, Alec Valenta, Xuefei Sun, Karl Mechtler

**Affiliations:** Thermo Fisher Scientific, Dornier Str. 4, 82110 Germering, Germany; IMP─Institute of Molecular Pathology, Campus-Vienna-Biocenter 1, A-1030 Vienna, Austria; Thermo Fisher Scientific, 1228 Titan Way, Sunnyvale, CA 94085, USA; IMBA─Institute of Molecular Biotechnology of the Austrian Academy of Sciences, Dr. Bohr Gasse 3, A-1030 Vienna, Austria; Gregor Mendel Institute of Molecular Plant Biology of the Austrian Academy of Sciences, Dr. Bohr Gasse 3, A-1030 Vienna, Austria

**Keywords:** bottom-up proteomics, single-cell proteomics, sample-limited proteomics, high sensitivity LCMS, nano LC-MS, UHPLC, Orbitrap, direct injection, trap-and-elute, data-independent acquisition, data-dependent acquisition

## Abstract

This study demonstrates how optimized liquid chromatography-mass spectrometry (LC-MS) conditions combined with a 50 µm internal diameter (I.D.) column operated at 100 nL/min enables high-sensitivity and high-throughput sample-limited proteomics analysis, including single-cell proteomics (SCP). Through systematic parameter evaluation, an optimized method was developed, capable of identifying ∼3,000 protein groups from 250 pg of HeLa protein digest using a 10-min gradient in the direct injection workflow using data-independent acquisition (DIA) from a library-free search method. Further improvements reduced the cycle time from 20 to 14.4 min by employing a trap-and-elute workflow, permitting 100 runs/day with 70% MS utilization. A proof of principle study indicated that *ca*. 1,700 protein groups were identified from single-cell samples without a library for label-free quantification (LFQ). In conclusion, we demonstrate a high-sensitivity LC-MS/MS configuration that serves the needs for limited sample analysis, permitting 100 runs/day throughout.

## Introduction

The past several years of proteomics have seen a significant shift toward proteome profiling of smaller and smaller sample quantities down to individual cells [1-4]. Obtaining high-quality data from sub-nanogram (ng) protein samples requires exquisite sensitivity, accuracy, and precision for all stages of the proteomics workflow, from sample collection to data analysis [5-9]. As such, state-of-the-art LC-MS technology plays a crucial role in our ability to investigate changes in protein expression without bias. While improvements in MS instrumentation have improved data collection speed and quality, mass spectrometers are still inherently limited by the quality of the sample at the LC-MS interface. High-quality, low-flow LC gradient separations provide minimal co-elution of target peptides, improving ionization efficiency (*i*.*e*., method sensitivity and, ultimately, proteome depth). Sensitivity is enhanced by utilizing ultra-low LC flow rates and small inner diameter columns/emitters [10-12], but at the potential cost of decreasing sample throughput in a single LC-MS configuration. Therefore, a balance between throughput and proteome depth is required for large-scale profiling of mass-limited samples. This is achieved through optimizing LC and MS parameters for fast sample injection and loading, efficient peptide separation, ionization and precursor isolation, accumulation, and fragmentation. Ideally, most method cycle time is dedicated to MS acquisition, with minimal time wasted on sample pickup and column equilibration. From this perspective, we optimized and evaluated the performance of a low-nanoflow UHPLC separation setup and several LC-MS methods in data-dependent acquisition (DDA), wide-window-acquisition (WWA) [13-14], and DIA to achieve deep proteome profiling at high throughput and high MS utilization for sample-limited proteomics, including single-cell analysis.

## Method

Full details on sample preparation, LC-MS conditions, and data analysis are provided in **Supplementary Method**.

### Sample preparation

Thermo Scientific™ Pierce™ HeLa Digest (A47996, 10 μg/vial) was reconstituted by adding 100 μL of 10% ACN (v/v) with 0.1% formic acid (FA) in water and sonicated for 5 min. Subsequently, a 10 μL sample was added to 990 μL water with 0.1% FA, after a few cycles of aspiration and release to get a concentration of 1 ng/μL HeLa digest in a 1 mL high recovery vial (6PSV9-V1), followed by either a 20 - 30s vertexing and 5 min sonication or a 20-min sonication prior to LCMS analysis. Sample amount was varied by injecting different volumes of HeLa digest onto the column.

With the label-free sample preparation workflow[7], the quality control (QC, prepared from 20 cells) and single-cell samples from HeLa and K562 cell lines (performance validation) were prepared in a 384-well plate and then covered by a silicon mat prior to LCMS analysis.

### Liquid chromatography parameters

The high-sensitivity low-nanoflow chromatography configuration comprises a Thermo Scientific™ Vanquish™ Neo UHPLC system (PN: VN-S10-A-01) and a Thermo Scientific™ Acclaim™ PepMap™ 100 C_18_ 50 µm I.D. × 15 cm column (PN: 164943) with a separation flow rate of 100 nL/min. The column inlet is connected to the sampler valve with a 10 µm x 350 mm (PN: 6250.5135) and 20 µm x 550 mm (PN: 6250.5260) Thermo Scientific™ nanoViper™ capillary via a low-dispersion Y-piece (PN:6250.1009) and a nanoViper blind nut (PN: 6040.2303, to block the third port) for direct injection and trap-and-elute workflows, respectively. The column outlet is connected to a 10 µm I.D. x 5 cm emitter (Fossillion technology, LOTUS) via a MicroTight unit (P771) or a PTFE sleeve (PN: 160489). A Sonation holder (PN: 004.800.01) insulated the Y-piece where the voltage was applied. The column was heated to 50 °C via a Sonation source (PN: PRSO-V2-ES72) with or without an inlay (PN: PRSO-V2-IZDV-72) for the MicroTight unit or PTFE sleeve, respectively. Mobile phase A and weak wash liquid were water with 0.1% FA (P/N LS118-500), and mobile phase B and strong wash liquid were 80% acetonitrile with 0.1% FA (P/N LS122500). All solvents were from Thermo Fisher Scientific. The autosampler temperature was 7 °C.

### MS parameters

The data was acquired on a Thermo Scientific™ Orbitrap Exploris™ 480 mass spectrometer with a Thermo Scientific™ FAIMS Pro™ interface. Three data acquisition strategies, DDA, WWA, and DIA, were employed to evaluate method sensitivity and performance.

### Data analysis

The LFQ-DDA dataset was processed with the Thermo Scientific™ Proteome Discoverer™ 2.5 software (version 2.5.0.400) using a 2-step SEQUEST™HT search algorithm and INFERYS™ rescoring node. The chimeric spectral in the LFQ-DDA and WWA-DDA datasets were searched with the CHIMERYS™ algorithm in Proteome Discover 3.0 (version 3.0.0.757), and DIA files were submitted to Spectronaut 17 (version 17.2.230208.55965) for peptide and protein identification and quantification. Only the identification by MS/MS spectrum was reported in the DDA dataset, and the false discovery rates (FDR) were all set below 1% at both the peptide and the protein levels. Further data analysis and plotting were performed with R script [13].

## Result and discussion

### A high-sensitivity and high-throughput configuration in direct injection workflow

Considering multiple factors for balancing the sensitivity (flow rate and column I.D.) and sample throughput (gradient delay volume, column pressure, and volume) in a low-flow LC-MS application, we established and evaluated the configuration using a 50 µm I.D. × 15 cm column in the direct injection workflow **(Figure 1A)**. Peptide ionization is carried out via a 10 µm I.D. glass emitter into a FAIMS pro interface operated at a single compensation voltage to reduce background ion interference. A liquid junction on the column inlet and zero-dead volume post-column connection to the emitter ensured stable ionization and minimal post-column dispersion to maintain chromatographic performance. Consequently, we developed five methods for fast sample loading at 1500 bar [16] (ca. 1 µL/min at 50 °C), efficient column washing, and equilibration in 2 min while maintaining a flow of 100 nL/min at a stable pressure. Each method required an additional 10 min for the sample injection cycle, column re-equilibration, and gradient delivery, enabling analysis of up to 72 samples/day **(Supplementary Figure 1)**.

**Figure 1.**
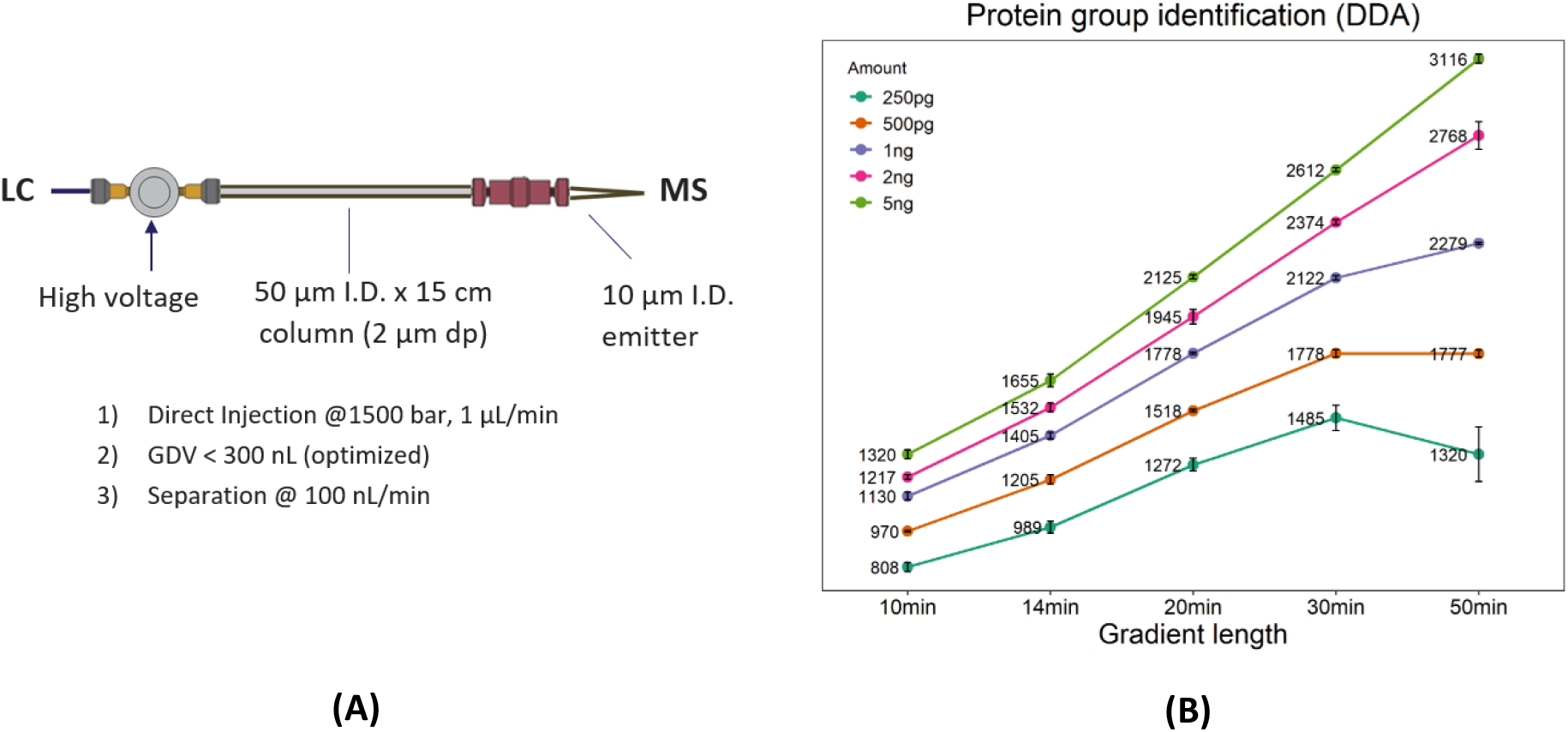
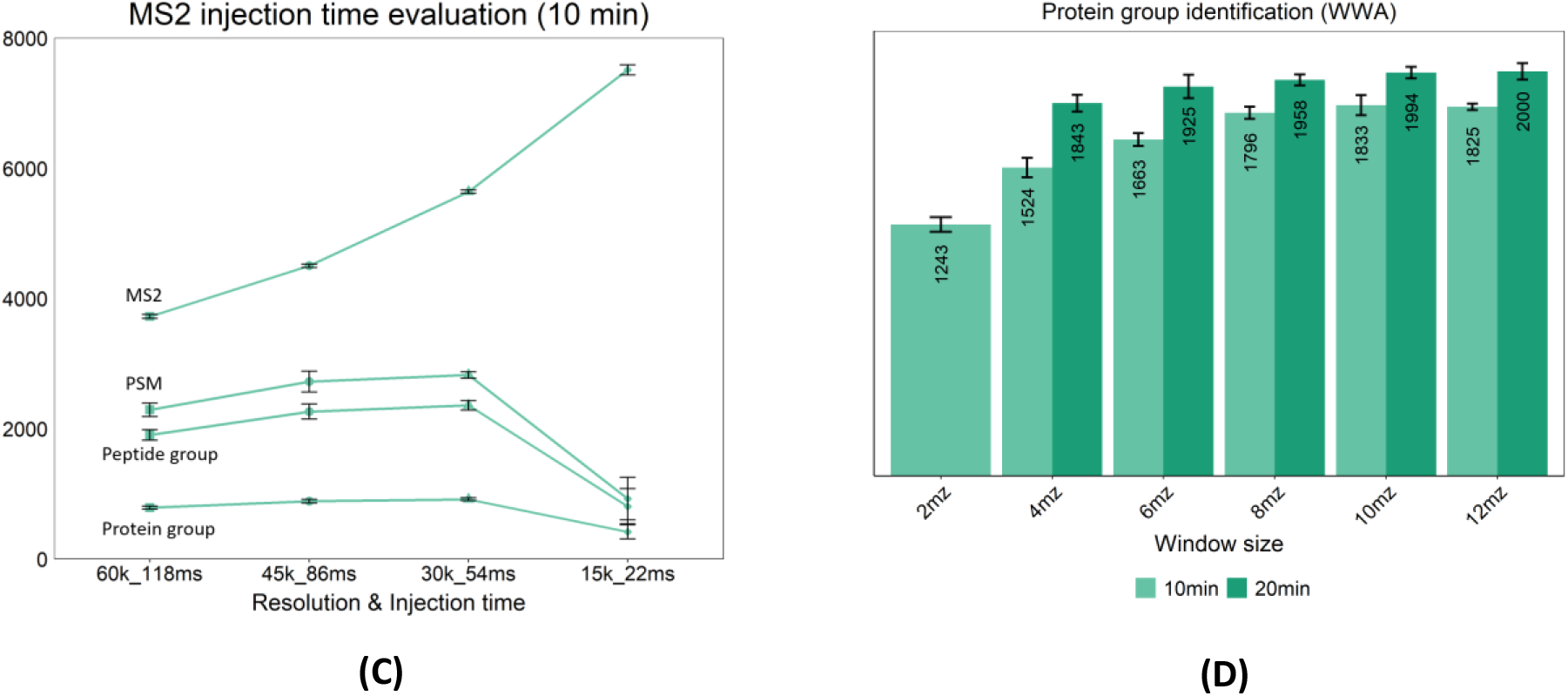
**(A)** Column configuration in direct injection workflow using 1500 bar for sample loading to increase sample throughput; **(B)** A linear increase of protein group identification from 250 pg to 5 ng HeLa digest; **(C)** The fastest scan speed using 22 ms MS2 IT impacted the I.D.s negatively with more MS2 not being translated into PSM, peptide and protein identifications; **(D)** 10/12 m/z isolation window significantly boost protein group identifications using WWA approach in a 10-min gradient

This configuration yielded < 4-second peak widths (median FWHM, only considering FWHM >0) and < 5 seconds retention time variation for the HeLa digest run using the 10-min gradient, regardless of the amount of sample injected (250 pg to 5 ng), *i*.*e*., the injected amount does not exceed the column loading capacity (**Supplementary Figure 2**). A linear increase in peptide and protein group identifications from 250 pg to 5 ng HeLa digest (**Figure 1B**) suggests excellent method suitability for analyzing limited sample amounts in the DDA mode. We identified ∼1,500 protein groups from 250 pg HeLa digest in a 30-min gradient without match-between-runs. To the authors’ knowledge, this represents the comprehensive DDA data with the most identifications to date using the conventional database search algorithm [17]. Moreover, ∼800 protein groups were identified using a 10-min gradient, suggesting the potential for WWA and DIA strategies to further boost proteome coverage by identifying/quantifying co-isolated peptides. Notably, peak broadening in the 50-min gradient reduced identifications for 250 and 500 pg samples, whereas injections with larger sample amounts (1, 2, and 5 ng) were unaffected due to more intense peaks and, consequently, more ions for sampling. Nevertheless, the low variation in protein abundance confirms reproducible LC-MS performance for protein quantification (**Supplementary Figure 3**), which is suitable for sample-limited proteomics, including single-cell proteomics (SCP).

### Enhance DDA performance by faster MS scan and chimeric spectrum deconvolution

Given the high sensitivity achieved in this configuration, we explored the effect of MS2 scan speed on DDA performance in a 10-min gradient with a 250 pg sample (single cell level amount) by decreasing the injection time (IT) from 118 to 22 milliseconds (ms), along with the corresponding Orbitrap resolution. Encouragingly, 10 -20% more peptide and protein identifications were gained from 54 ms IT and 30,000 resolution. This results from the high-sensitivity front end (stable low-nano flow rate over the gradient, high ionization efficiency, and background ion filtering by the FAIMS Pro interface) providing sufficient ion clusters/time for faster scanning (**Figure 1C**). However, IT below 54 ms had the opposite impact on identifications, where more MS2 scans did not translate into more peptide spectrum matches (PSMs) and peptide and protein Identifications. It illustrates that ion accumulation time and spectral resolution are crucial for precursor fragmentation and spectrum identification to analyze limited sample amounts.

Precursor co-isolation in DDA mode commonly results in complex spectra that limit the performance of conventional database search algorithms, e.g., SEQUEST, for peptide identification. By employing CHIMERYS, an advanced AI-driven algorithm for spectral deconvolution, we boosted protein Identifications for 250 pg and 5 ng (10-min gradient) by 40% (1142) and 80% (2360), respectively, from the same dataset. However, the longer, 30-min gradient only gained 8% more protein Identifications due to better chromatographic resolution between peptide peaks, yielding less complex MS spectra (**Supplementary Figure 4)**. Further exploration with a WWA strategy in MS1 combined with CHIMERYS enabled 1,800 protein group Identifications (an increase of 120%) from 250 pg HeLa digest with a 10-min gradient and 10 Th isolation window (**Figure 1D**). This result agrees with our previous findings using a different LC-MS configuration [7].

### Achieve next-level performance with DIA

Since DIA can sample more ion population clusters for fragmentation and has seen recent advances in library-free identification algorithms, a systematic evaluation of different window sizes in DIA was performed. It yielded an average of as much as 3,000 protein group identifications from a 250 pg HeLa digest using a 10-min LC gradient applying a 20 m/z isolation window **(Figure 2A)**. A decreasing identification trend illustrates the challenge of deconvoluting extraordinarily complex spectra using wider isolation windows. Nevertheless, the DIA approach enables deep proteome coverage with low quantification variation across > 4 orders of magnitude dynamic range **(Supplementary Figure 5)**. Balancing the required data points for protein quantification and proteome depth, we employed a 40 m/z isolation window further to explore the potential of this short gradient method. An increasing trend in protein identification was seen with increased sample amount where >4,600 protein groups were identified, of which 87% were quantified at <20% CV covering > 4 orders of magnitude of dynamic range from 10 ng sample **(Figure 2B and 2C)**.

**Figure 2.**
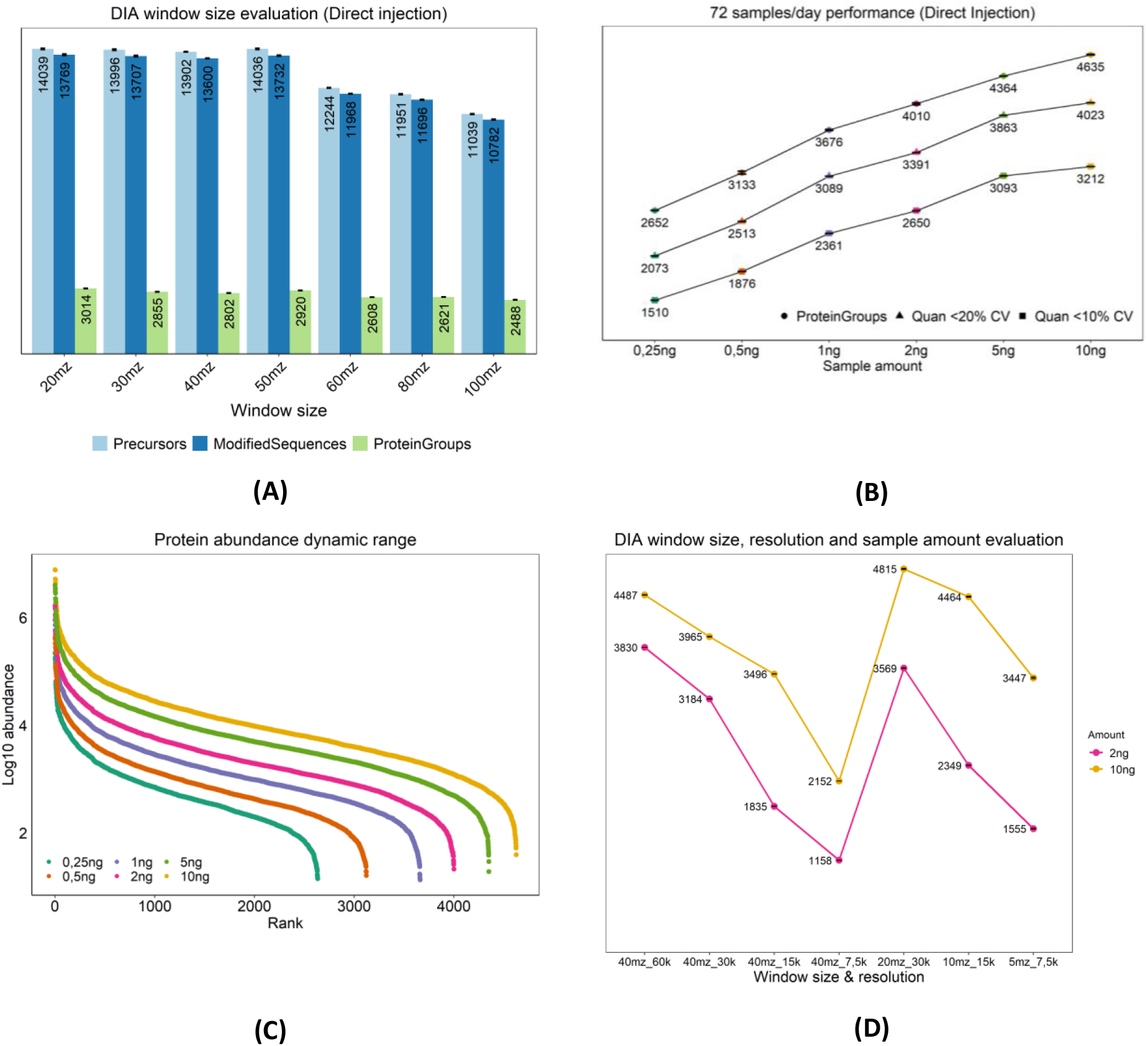
**(A)** More than 3,000 proteins groups were identified on average from 250 pg HeLa sample using a 10-min LC gradient; **(B)** A linear increase of protein group identification from 250 pg to 10 ng HeLa digest in DIA; **(C)** The protein abundances span more than 4 orders of magnitude of the dynamic range; **(D)** DIA with a wider window requires a higher resolution MS2 for spectral deconvolution but still shows dependency to sample amount

Due to the high sensitivity achieved from the sub-nano gram samples, we further explored the relationship of the MS2 IT, resolution, MS1 window size, and injection amount for larger sample quantities (**Figure 2D**). In short, reducing the resolution and corresponding IT (*i*.*e*., increasing scan speed) decreased protein Identifications and quantification for 2 and 10 ng samples, illustrating that the MS2 complexity in this condition requires 60,000 resolutions for spectral deconvolution. While keeping scan speed, the window size must be coordinated with the sample amount to reflect spectral complexity and ion intensity to enable optimal protein identification. For instance, a window size of 20 m/z would increase identifications by 10% for a 10 ng sample using an MS resolution of 30,000. Moreover, a longer gradient of 20-min does not significantly increase peptide and protein identifications, indicating that a 10-min gradient is optimal for DIA-based sample-limited proteomics **(Supplementary Figure 6)**.

### Increased sample throughput with a trap-and-elute workflow for LFQ-DIA SCP profiling

To accelerate sample loading and eliminate the potential negative impact of impurities and detergent on electrospray ionization, we employed a trap column operated in a backward flush mode to maintain peak shape, successfully decreasing the method cycle time to 14.4 min (100 samples/day) for a 10-min gradient at 100 nL/min with approximately 70% MS utilization (**Figure 3A**). Using the optimized parameters from the direct injection workflow, we observed ca. 14% - 17% lower protein identifications in the trap-and-elute workflow (**Figure 2B** & **3B**). Nevertheless, with the performance of over 2,200 identified protein groups in 250 pg >1,100 protein groups were still identified from as low as 60 pg samples (**Figure 3B**). Consequently, this configuration shows a similar sample throughput and MS utilization as a remarkable dual-trap fluidic configuration but at a five times lower flow rate using a column with a 1.5 times narrower I.D., achieving higher ionization efficiency and sensitivity [18]. Ultimately, this method identified >1,200 protein groups from HeLa and K562 QC samples in LFQ-DIA with 100 cells/day (CPD) throughput (**Figure 3C**), regardless of the potential sample loss resulting from the manual transfer step, long-distance transportation and freeze-thaw cycle [7]. Method performance was reproducible when analyzing QC samples from the same batch between two labs **(Figure 3C)**, regardless of different operators, columns, emitters, LC, MS *etc*. A further step of work illustrated the high performance of the method by identifying > 1,700 protein groups in HeLa cells with library-free DIA **(Figure 3D)**. Notably, it outperformed our benchmarked dataset **(Figure 3D) [7]** with 20% more identified protein groups with three times higher sample throughput, and the chromatography configuration is also comparable with different MS settings (**Supplementary Figure 7)** [19].

**Figure 3.**
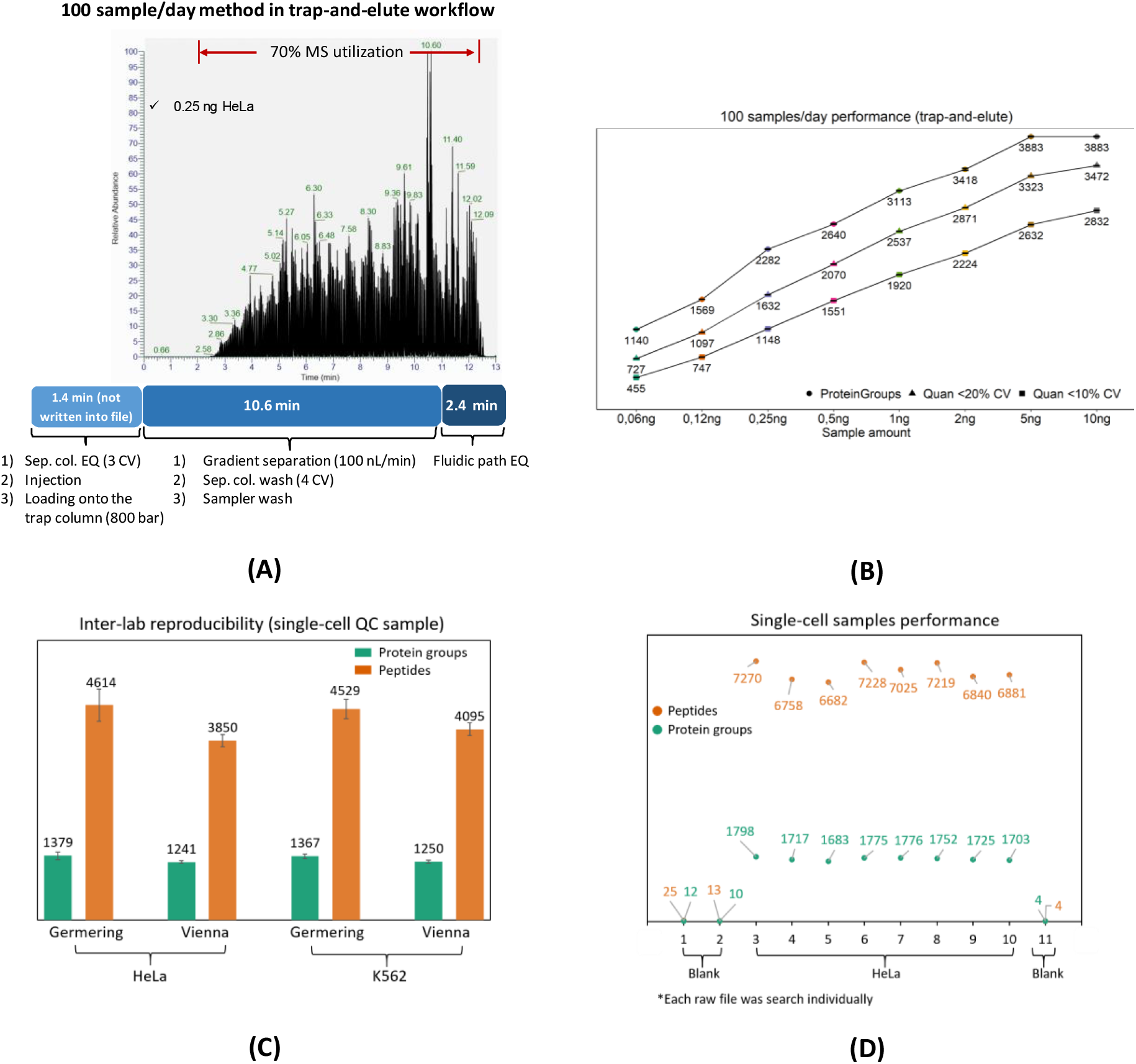
**(A)** 14.4-min method in trap-and-elute workflow permits 100 samples/day throughput at 100 nL/min; **(B)** A linear increase of protein group identification from 60 pg to 10 ng HeLa digest in DIA; **(C)** Reproducible inter-lab performance verifies the high-sensitivity of the method in HeLa and K562 QC samples; **(D)** Achieving *ca*. 1,700 protein groups identification from single HeLa cell with neglectable carryover

## Conclusion

We established a high-performance configuration for sample-limited proteomics and validated it with single-cell samples after systematically evaluating the impact of sample amount, gradient length, MS2 IT, and database search algorithms. It utilizes a 50 µm I.D. separation column at 100 nL/min to achieve high-sensitivity results where *ca*. 1,700 protein groups are identified from HeLa single cells at a throughput of 100 CPD, which outperformed our previously published results from a 32 CPD LC-MS method with same sample preparation workflow. It is an excellent platform meeting the needs of high sensitivity in limited sample analysis, *e*.*g*., SCP and spatial Omics, for investigating cellular heterogeneity in clinically and biologically relevant cellular populations.

## Acknowledgments

This work was funded by the EPIC-XS, Project Number 823839, the Horizon 2020 Program of the European Union, project LS20-079 of the Vienna Science and Technology Fund, the Era-Caps I 3686-B25, and the project P35045-B of the Austrian Science Fund. All LC-MS/MS analyses in Vienna were performed on instruments of the Vienna BioCenter Core Facilities instrument pool. We thank Ulla Schellhaas at IMP for providing us with K562 cells.

Furthermore, special gratitude goes to Alexander Makarov, Alexander Boychenko, Christopher Pynn, Wim Decrop, Martin Samonig, Cornelia Boeser, Jeff Op de Beeck, Tabiwang Arrey, and Dominic Hoch from Thermo Fisher Scientific for the technical support and fruitful discussion.

