## Supplementary Figures for "A high-sensitivity low-nanoflow LC-MS configuration for high-throughput sample-limited proteomics"

### A high-sensitivity low nano-flow chromatography configuration for high-throughput MS-based sample-limited proteomics

| Sample Throughput/Day | Cycle Time (min) | Gradient Length (min) | Elution Window (min) | Sample Injection & Loading (min)* | MS utilization (%) |
|---|---|---|---|---|---|
| 72 | 20 | 10 | 11 | 5.4 | 55 |
| 60 | 24 | 14 | 14 | 5.4 | 62.5 |
| 48 | 30 | 20 | 21 | 5.4 | 70 |
| 36 | 40 | 30 | 31 | 5.4 | 77.5 |
| 24 | 60 | 50 | 51 | 5.4 | 85 |

* 1 µL sample injection + 2 µL loading

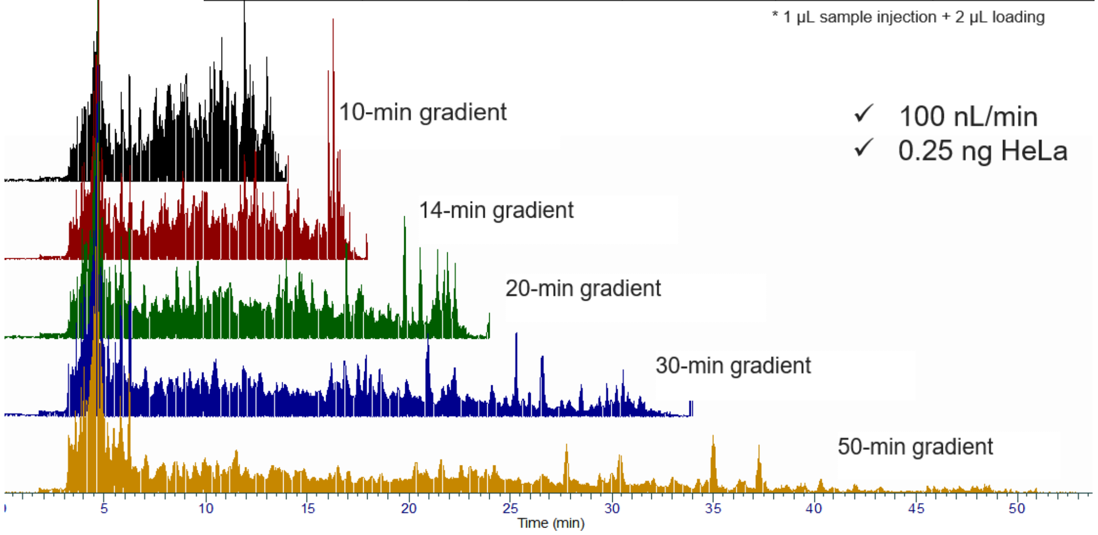


**Supplemetary Figure 1.** Five optimized direct injection LC-MS methods balancing sensitivity and throughput for different proteome coverage needs

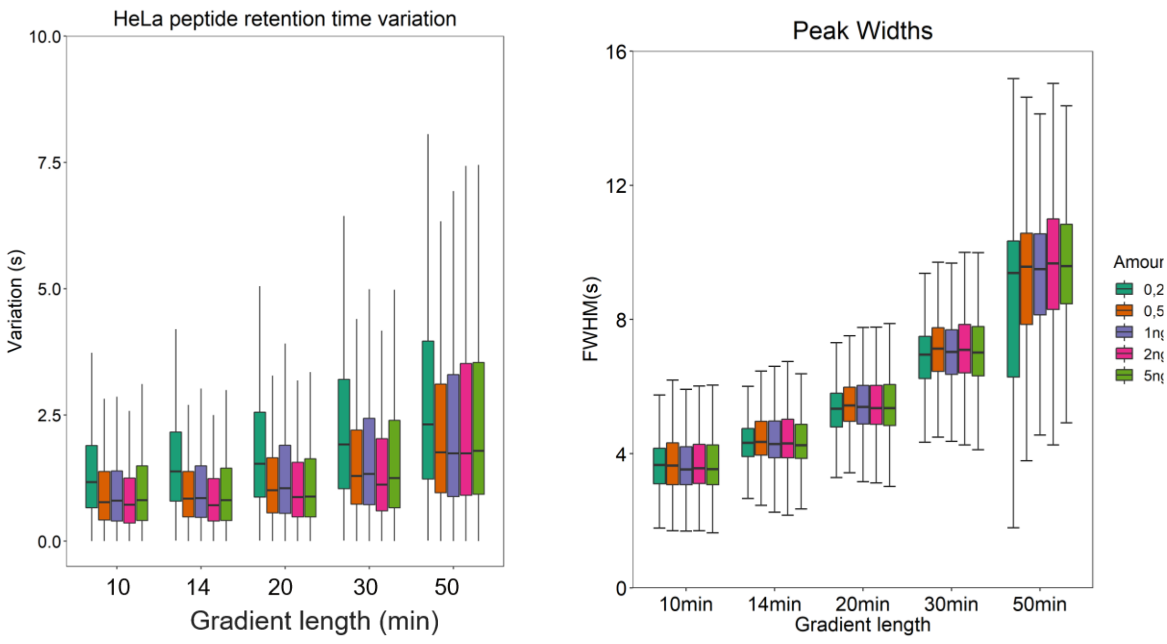


(A) (B)

**Supplemenetary Figure 2**. Low retention time variation (A) and narrow peak width (B)

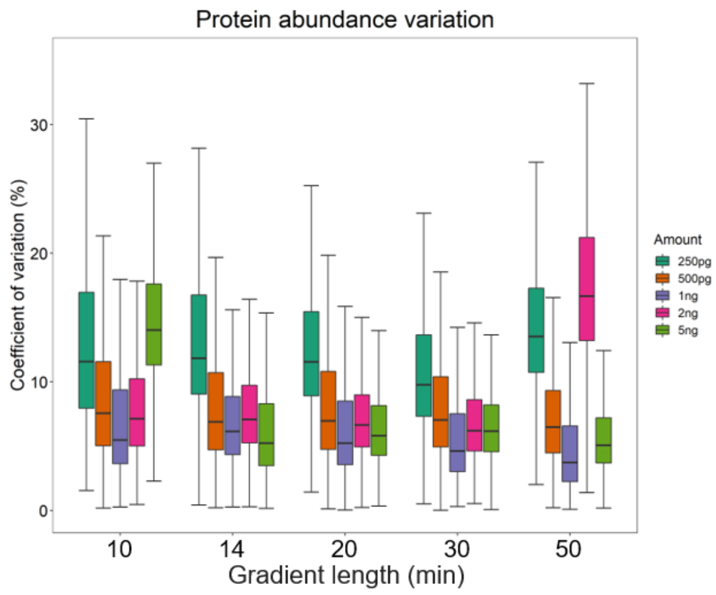


**Supplemenetary Figure 3**. high quantitation accuracy of the LC-MS DDA performance

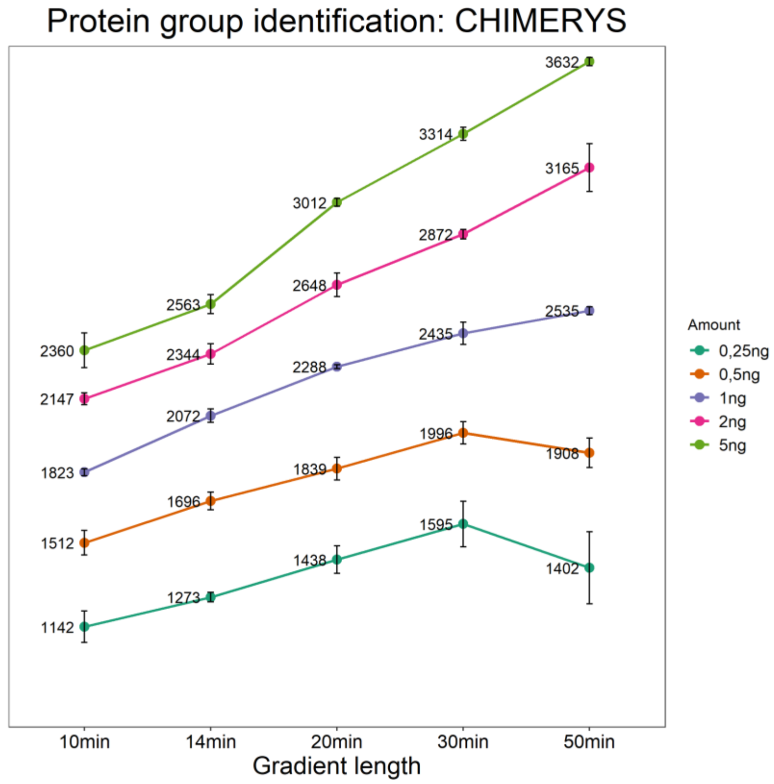


**Supplemenetary Figure 4**. Improved protein group identification by CHIMERY algorithm

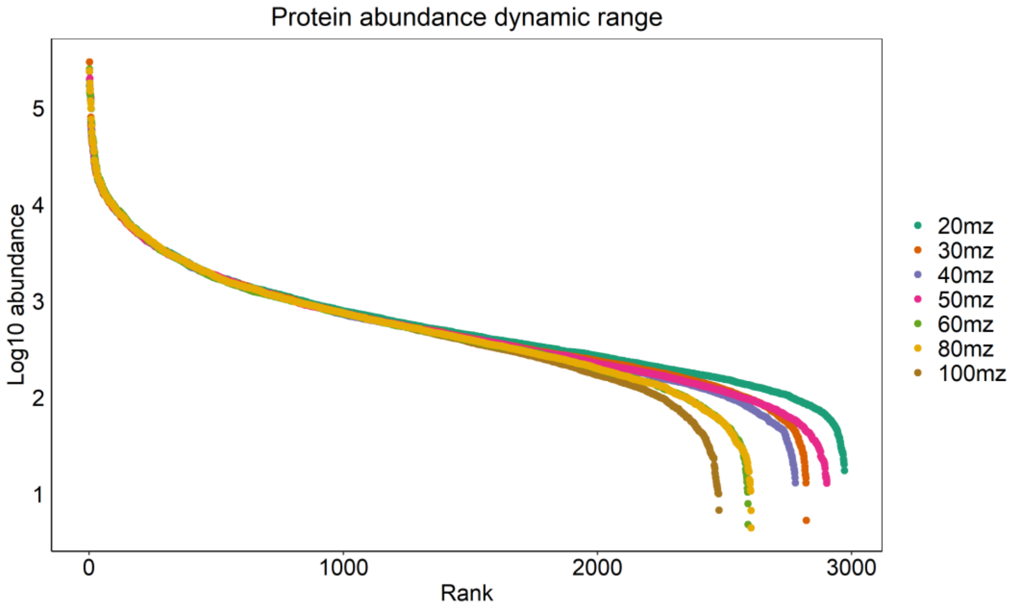


**Supplemenetary Figure 5**. Protein dynamic range spanning 4 orders of magnitude with 250 pg samples in 10-min gradient

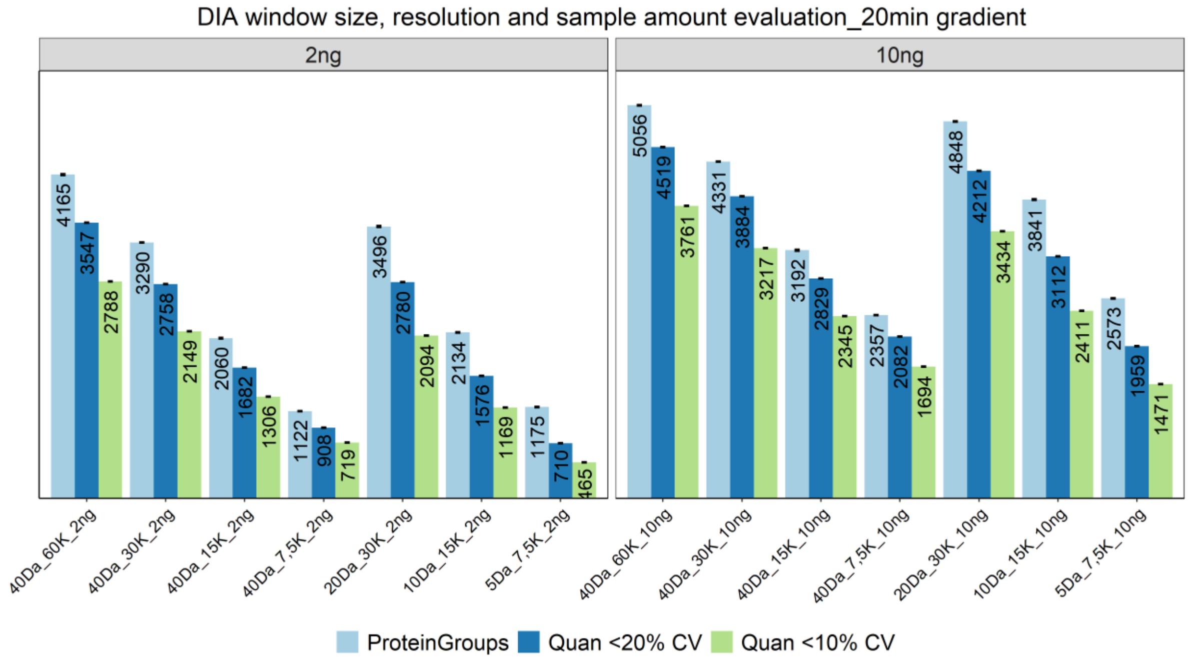


**Supplementary Figure 6**. Direct injection workflow performance in a 20-min gradient

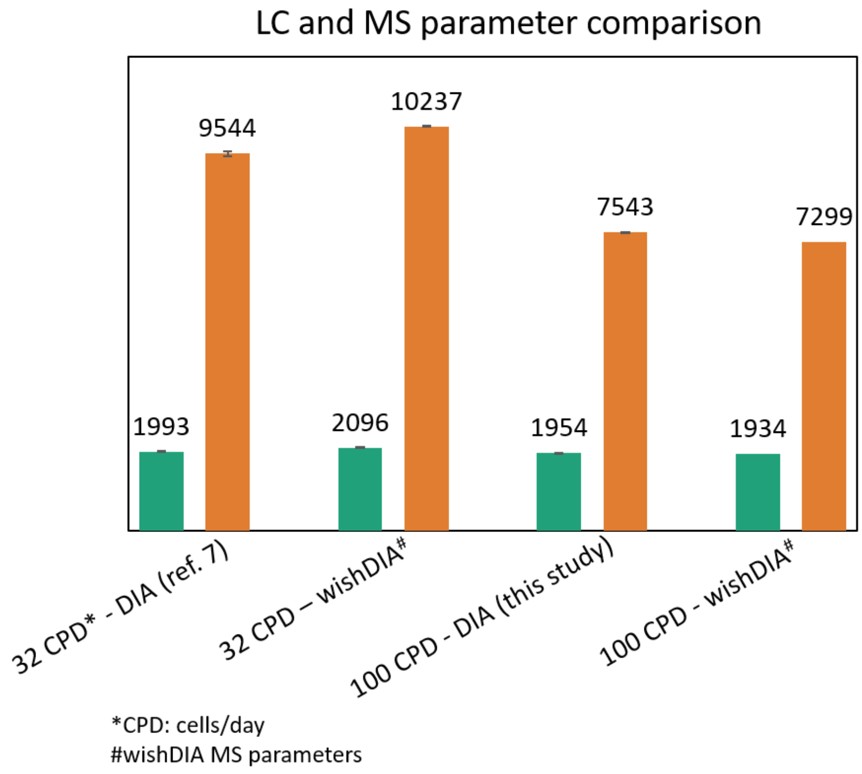


**Supplementary Figure 7**. Reproducing benchmarked performance with trap-and-elute workflow for LFQ-DIA analysis at 100 CPD throughput
